# Single-cell analysis of childhood leukemia reveals a link between developmental states and ribosomal protein expression as a source of intra-individual heterogeneity

**DOI:** 10.1101/683854

**Authors:** Maxime Caron, Pascal St-Onge, Thomas Sontag, Yu Chang Wang, Chantal Richer, Ioannis Ragoussis, Daniel Sinnett, Guillaume Bourque

## Abstract

Childhood acute lymphoblastic leukemia (cALL) is the most common pediatric cancer. It is characterized by bone marrow lymphoid precursors that acquire genetic alterations, resulting in disrupted maturation and uncontrollable proliferation. More than a dozen molecular subtypes of variable severity can be used to classify cALL cases. Modern therapy protocols currently cure 85-90% of cases, but other patients are refractory or will relapse and eventually succumb to their disease. To better understand these difficult cases, we investigated the nature and extent of intra-individual transcriptional heterogeneity of cALL at the cellular level by sequencing the transcriptomes of 39,375 individual cells in eight patients (six pre-B and two pre-T) and three healthy pediatric controls. We observed intra-individual transcriptional clusters in five out of the eight patients. Using pseudotime maturation trajectories of healthy B and T cells, we obtained the predicted developmental state of each leukemia cell and observed distribution shifts within patients. We showed that the predicted developmental states of these cancer cells are inversely correlated with ribosomal protein expression levels, which could be a common contributor to intra-individual heterogeneity in cALL patients.

## Introduction

Childhood acute lymphoblastic leukemia (cALL) is the most frequent pediatric cancer, accounting for ∼25% of all pediatric tumors. This cancer is characterized by bone marrow lymphoid precursors that acquire sequential genetic alterations, resulting in disrupted maturation and uncontrollable proliferation. Precursor B cell ALL (pre-B ALL) represent ∼85% of cases and precursor T cell ALL (pre-T ALL) ∼15%, which can be further subdivided into more than a dozen molecular subtypes. The high hyper diploid cases (HHD) and those harboring the t(12;21) [*ETV6/RUNX1*] rearrangement represent about ∼60% of pre-B cALL cases and are associated with a good prognosis^1,2^. Other less frequent (< 10%) subtypes, such as MLL-rearranged, t(9;22) [*BCR/ABL1*] or pre-T are associated with intermediate to poor outcomes^1,2^. Five-year event-free survival rates have seen remarkable progress since the 1960s and are now up to ∼85-90% thanks to patient risk stratification and effective treatment combinations^3^. Despite this therapeutic success refractory and relapsing cALL still represents one of the most frequent cause of death by disease in pediatrics^4^. The disease follows a two-hit model starting with initiating genetic lesions in lymphoid progenitors (genomic translocations, copy number alterations, point mutations) occurring in utero or very early in life and subsequent secondary genetic lesions driving the leukemogenesis^3^.

Somatic mutations at diagnosis arise in many genes involved in multiple biological pathways such as B cell development (e.g. *PAX5, IZKF1*), signaling (e.g. *RAS, NRAS, CRLF2*) epigenetic regulation (e.g. *CREBBP, KMT2D*) and cell cycling (e.g. *CDKN2A*)^5^. This led to the proposition of clonal evolution characterized by a branching model in which leukemic subclones acquire sequential genetic alterations conveying fitness advantages^5^. In a significant number of relapse cases, ancestral minor subclones present at diagnosis become resistant and positively selected under treatment^5,6^. Next-generation sequencing (NGS) has recently provided meaningful insights toward the characterization of somatic events providing selective advantages to tumor cells. Mutations in RAS pathway genes have been associated with early relapse and chemoresistance in cALL whereas mutations in the cytosolic 5’-nucleotidase II gene (*NT5C2*) were shown to be involved in resistance to treatment^7,8^. A study on the rise and fall of subclones revealed that the majority of patients harbor at least one genetic subclone at diagnosis and three quarters of relapse cases were from resistant minor ancestral subclones^5^. Another study found the presence of at least two distinct genetic subclones in five out of six patients using targeted single-cell DNA sequencing^9^. In this context, we reported different clonal dynamics between early (during treatment) and late (>36 months after diagnostic) bone marrow relapse events and showed variable clonal origin in late relapse events^10^.

Genetic alterations of subclones can provide information on the perturbed functions of genes and their corresponding pathways but the transcriptional impacts of these alterations in cALL are unknown as intra-individual transcriptional heterogeneity is not well characterized. Additionally, subclonal epigenetic modifications could cause changes in gene expression that cannot be captured by looking at genetic alterations only. Transcriptional changes at the single cell level can be assessed using recent single cell technologies that enable the epigenetic profiling of thousands of single cells, including droplet-based whole transcriptome profiling (scRNA-seq)^11^. Profiling the transcriptomes of single cells in both diagnosis and relapse samples could help identify expression signatures of relapsing subclones in hematological cancers^12^. The prognostic value of scRNA-seq was previously demonstrated in a single cell gene expression study that assessed the transcriptional heterogeneity of patients initially assigned to a unique subtype that were further classified into additional subgroups with distinct survival^13^. In this study we sought to uncover the extent and nature of transcriptional heterogeneity in cALL patients at diagnosis, identify distinct subpopulations of cancer cells and uncover deregulated genes and pathways. These intra-individual transcriptional differences have not been characterized and could provide information on the impact of genetic or epigenetic alterations on the expression programs of these cells and help us gain additional knowledge on the mechanisms of disease development and relapse.

## Results

We generated single cell gene expression data from 39,375 pediatric bone marrow mononuclear cells (BMMCs) from eight cALL patients of common subtypes: 4 ETV6-RUNX1, 2 High Hyper Diploid (HHD) and 2 pre-T (> 50% blasts) and 3 healthy donors (Table 1). We assessed the quality of our data in healthy pediatric BMMCs with available data in healthy adult BMMCs (10X Genomics, Methods). We recovered the expected cell types using known cell surface marker genes (Figure 1A-D). In healthy cells, the predicted cell cycle phases showed a higher proportion of cycling cells in B cells and Immature erythrocytes than in other cell types (Figure 1E). By combining healthy pediatric BMMCs with cALL cells (n=38,922 after quality control), we observed distinct clusters of healthy and cancer cells (Figure 2A). Healthy clusters were determined using both healthy cell cluster assignment (Supp. Figure 1A) and expression of cell surface marker genes. Between 2 and 60% of cALL cells per sample clustered with healthy pediatric BMMCs of different cell types (Figure 2C, Supp. Figure 1B). These cells likely represent non-cancerous cells normally found in samples of variable tumor purity (due to disease severity or technical variability), rather than lineage switching cancer cells or cancer cells having healthy-like transcriptional profiles. When looking at the predicted cell cycle phases of cALL cells, we observed a continuous spectrum of phases G1→S→G2/M on the UMAP representation (Figure 2B). For six out of eight patients, cells were mostly in the G1 phase (Figure 2D). Many methods can correct for different sources of transcriptional variation^14,15^, however regressing out the cell cycle phase in our data failed to completely remove this effect as we could still observe clusters of cells in cycling phases on UMAP (Supp. Figure 1C). Thus, in further analysis, we decided to reduce the expression variability by keeping cancer cells that did not cluster with healthy cells (remaining n=25,788; ∼79.5%) and that were in G1 phase only (remaining n=16,731; ∼51.6%; Figure 3A). We looked at the transcriptional profiles of these cancer cells using non-supervised hierarchical clustering of the hundred most variable genes. We observed two distinct clusters of pre-B and pre-T cells as shown by the expression of the *CD79A/B* and *CD3D* surface markers (Figure 2E, Supp. Figure 1 D-F). We also noticed transcriptional differences between pre-B subtypes (e.g. *RAG1* over expressed in ETV6-RUNX1 samples) and individuals (e.g. *TCL1B* over expressed in samples ETV6.RUNX1.3/ETV6.RUNX1.4 and *CD1E* over expressed in sample PRE-T.1). These transcriptional profiles corroborate expected cell surface marker expression for these types of cells and indicate good quality of the single cell expression data.

**Figure 1.**
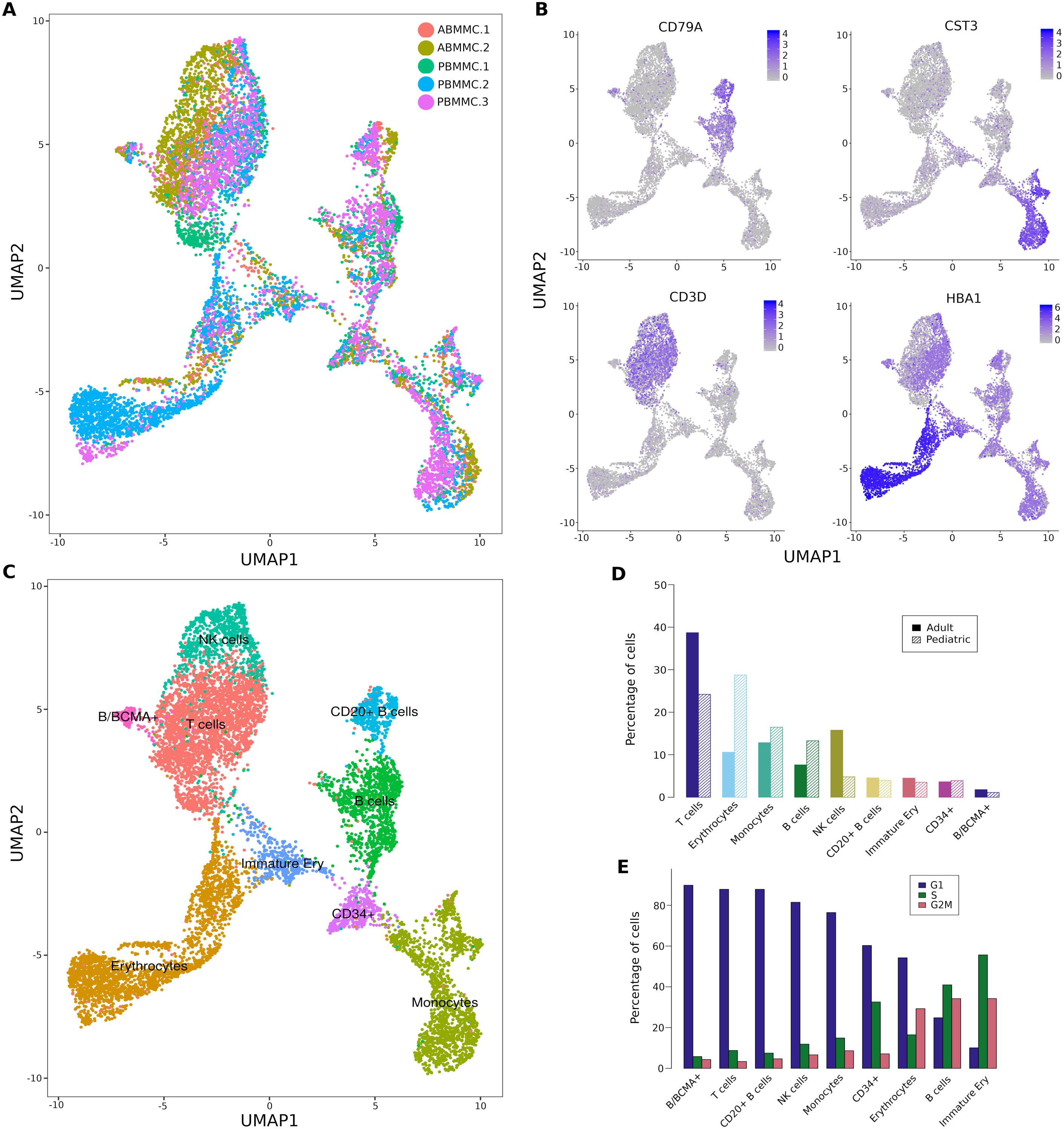
Cell types identified in healthy pediatric and adult bone marrow mononuclear cells (BMMCs) using single cell RNA-seq. **A)** UMAP representation of healthy BMMCs from three pediatric (PBMMC; n=6,836 cells) and two adult (ABMMC; n=3,467 cells) donors. **B)** Expression of cell surface marker genes used to assign cell types: *CD79A* (B cells), *CST3* (Monocytes), *CD3D* (T cells) and *HBA1* (Erythrocytes). **C)** Cell types identified in healthy pediatric and adult BMMCs. **D)** Proportion of cells of a given cell type in pediatric and adult BMMCs. **E)** Proportion of healthy cells in predicted cell cycle phases per cell type (G1, S and G2/M).

**Figure 2.**
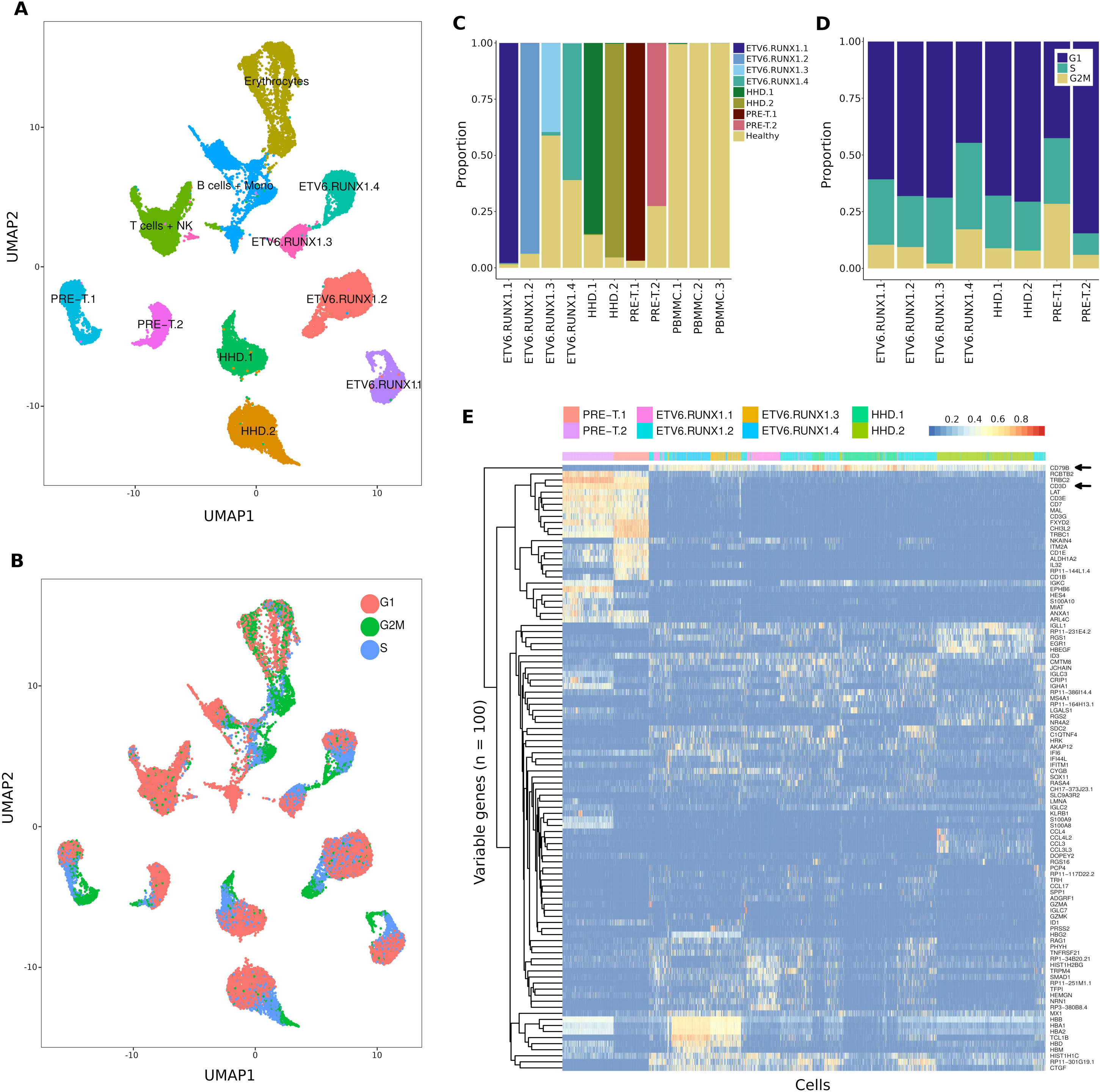
Transcriptional landscape of cALL cancer cells. **A)** UMAP representation of BMMCs from three healthy pediatric donors (n=6,836 cells) and eight cALL patients (n=32,086 cells). **B)** UMAP representation of predicted cell cycle phases for healthy and cancer BMMCs. **C)** Proportion of cells clustering with healthy cell clusters. **D)** Proportion of cancer cells in predicted cell cycle phases (G1, S and G2/M). **E)** Heatmap and unsupervised clustering of normalized and scaled expression of the top 100 most variable genes in leukemia cells.

**Figure 3.**
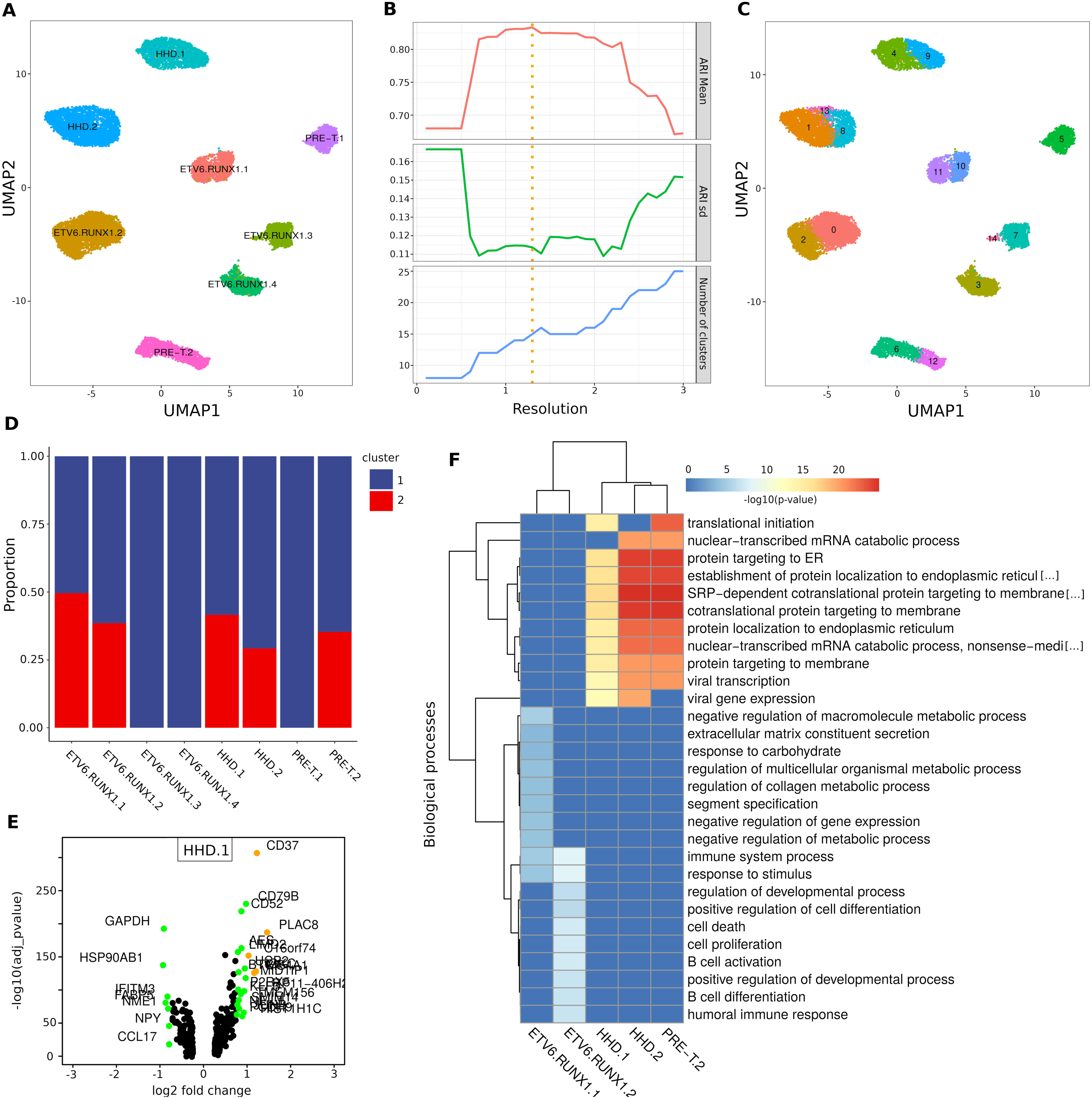
Intra-individual transcriptional heterogeneity reveals deregulated genes and pathways within cALL samples. **A)** UMAP representation of cALL cells in G1 phase not clustering with healthy cell clusters (n=16,731). **B)** Mean Adjusted Rand Index (ARI) of clustering solutions over a range of resolutions (highest mean ARI at 1.3 resolution). **C)** Clusters of cells identified in cALL samples using the highest mean ARI resolution. **D)** Proportion of cells belonging to each intra-individual cluster after removing clusters having less than 10% of cells (n=16,162). **E)** Differentially expressed genes between two the clusters of cells within the HHD.1 sample (log fold-change > 0.75 = green, > 1 = orange). **F)** Heatmap and unsupervised clustering of enriched GO biological pathways obtained using the top 100 most significant differentially expressed genes of cALL samples.

### Intra-individual transcriptional heterogeneity

We assessed the intra-individual transcriptional heterogeneity by identifying expression profile signatures unique to subsets of cancer cells within a sample. These intra-individual transcriptional differences could highlight deregulated genes in potentially resistant subclones or reveal epigenetic changes driven by subclonal genetic alterations. We applied a strategy that would return the most representative clustering solution over a range of clustering resolutions. Multiple clustering solutions were generated over a range of parameters and pairwise Adjusted Rand Index (ARI) values were computed (Figure 3B). We retained the clustering solution that had the highest mean ARI and identified at least two clusters in six samples, but no intra-individual transcriptional clusters for samples ETV6.RUNX1.4 and PRE-T.2 (Figure 3C). Transcriptional differences between subsets of cell within a sample require enough cells (>100) to be properly powered for differential expression analyses^16^. Thus, we discarded clusters with less than 10% of total cells for samples ETV6.RUNX1.3 and HHD.2 (remaining n=16,162). This filtering steps resulted in 5 samples with two transcriptional clusters with proportions of cells ranging from ∼25 to 50% (Figure 3D). We performed intra-individual differential gene expression analyses for the two subsets of cells within these 5 samples. Genes related to B and T cell maturation (e.g. *CD37, CD79B, JCHAIN, IGLL1, VPREB3, CD52*), ribosomal protein genes (*RPS-*, RPL-\**) and cancer/stress genes (e.g. *PRAME, ARHGDIB, FOS, JUN*) (Figure 3E, Supp. Table 3) were within the most significantly deregulated genes between clusters in those samples. Gene ontology analyses on the five differential expression gene lists led to a more general understanding of the modulated biological pathways within each sample (Supp. Table 4). The samples were then clustered based on the top ten most significantly enriched biological pathways. We observed two distinct clusters: one cluster of ETV6-RUNX1 samples showing modulated pathways in B cell activation/differentiation, cell proliferation/death and regulation of expression and metabolic processes and another cluster of HHD and pre-T samples showing modulated pathways in translation initiation and protein synthesis (Figure 3F). These results demonstrate that we can detect distinct sources of transcriptional variation between subsets of cells within a sample.

### Developmental states and ribosomal protein gene expression

To address whether the observed intra-individual transcriptional variability could be linked to the maturation and differentiation states of the cancer cells, we implemented two developmental state classifiers based on the expression profiles of healthy B and T cells. We assigned pseudotime values of 0 to 1 along the maturation spectrum from stem-like to mature developmental states (Figure 4A). The performances of the classifiers were assessed on healthy cells using cross-validation and showed good predictive performance using average Root Mean Squared Error (RMSE) (Figure 4B). Final classifiers were trained on all healthy B and T cells and then used to obtain the predicted developmental states of every cancer cell. We observed statistically significant shifts in developmental pseudotime distributions within samples (Figure 4C). The most significant shifts were found within HHD and PRE-T samples, while less pronounced shifts were observed within ETV6-RUNX1 samples. These findings correlate with differential expression results that revealed modulated expression of ribosomal protein genes in HHD and PRE-T samples (Supp. Table 3). An inverse relationship between maturation states of normal hematopoietic cells and ribosomal protein (RP) expression levels was previously reported in a zebrafish single cell RNA-seq study^17^. We observed the same relationship for cALL cancer cells in all samples exhibiting intra-individual transcriptional clusters where subsets of cells with higher pseudotime had lower ribosomal protein expression levels (Figure 4D). This trend was clearer in samples having strong (e.g. HHD.1) vs weak (e.g. ETV6.RUNX1.2) shifts in pseudotime (Figure 4E). Ordering cells by RP expression revealed a gradient of expression that was correlated to clusters having similar behavior (Figure 4F). These findings suggest that developmental states and RP expression could contribute to the intra-individual transcriptional heterogeneity observed in some cALL patients.

**Figure 4.**
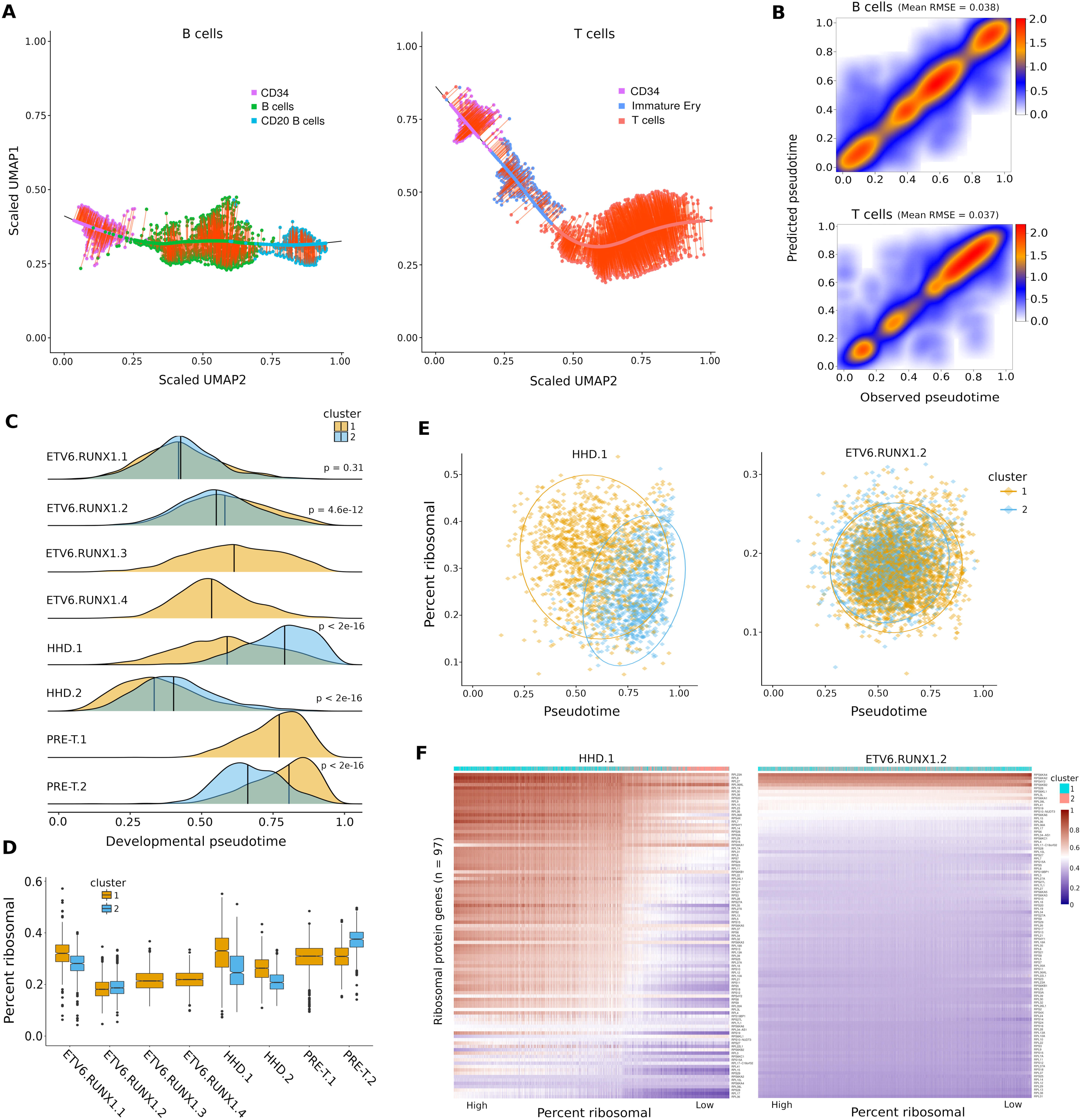
Predicted developmental state is inversely correlated with ribosomal protein expression in cALL cells and is a major source of intra-individual transcriptional heterogeneity. **A)** Left: UMAP representation of the maturation spectrum of healthy pediatric and adult B cells (CD34+ →B cells → CD20+ B cells) used for the B cell developmental state classifier. Right: UMAP representation of the maturation spectrum of healthy pediatric and adult T cells (CD34+ → Immature Erythrocytes →T cells) used for the T cell developmental state classifier. Cells were projected onto the loess fit of the spectrum and assigned a pseudotime value of 0 to 1 from the first stem-like cell to the last mature cell. **B)** Observed vs predicted pseudotime of healthy B and T cells using a hundred 70/30 cross-validation splits; mean RMSE was computed over all splits. **C)** Density of predicted developmental state pseudotime distributions of leukemia cells per sample and intra-individual transcriptional cluster. **D)** Boxplot of ribosomal protein (RP) expression percentage in leukemia cells per sample and intra-individual transcriptional cluster. **E)** Pseudotime vs ribosomal protein expression in samples showing strong (HHD.1) vs weak (ETV6.RUNX1.2) intra-individual pseudotime and ribosomal protein expression shifts. **F)** Heatmap of normalized and scaled expression of ribosomal protein genes per cell sorted from high to low. An expression gradient correlated to cluster assignment can be observed in sample HHD.1 but not ETV6.RUNX1.2.

### Genetic alterations and transcriptional variability

To uncover genetic alterations that could be linked to intra-individual transcriptional heterogeneity, we first looked whether large cluster specific copy number variants (CNVs) could be linked to transcriptional clusters. We generated thirty equal size ‘metacells’ for each cluster to reduce noise and called copy number events using healthy pediatric BMMCs as controls. We recovered most copy number events seen with exome sequencing data and identified cluster specific copy number events (Figure 5A, Supp. Figure 3). We observed cluster specific copy number gains on chromosomes 21 and 22 in sample ETV6.RUNX1.1 and deletions of chromosome 5 and the p arm of chromosome 12 in sample ETV6.RUNX1.2. Subclonal copy number depth ratios were also seen for some these alterations using exome data (Supp. Figure 4). We did not observe large cluster specific copy number events for HHD or pre-T samples. We then looked whether there was a correlation between the presence of genetic subclones and intra-individual transcriptional clusters. We used somatic mutations from matched normal and tumor exome data (Supp. Table 5) and found no major differences in both the number of mutations and genomic locations between samples (except for sample ETV6.RUNX1.1 that had less mutations) (Figure 5B, C). We identified known cALL mutations in signaling proto-oncogenes (*ABL1, ROS1, NOTCH1*), cell cycle/epigenetic regulators (*CREBBP, CDKN2A*) and noticed few overlapping non-synonymous exonic mutations between samples (Figure 5D). We used all sufficiently covered (> 100X) somatic mutations to generate genetic evolution models and at least one minor genetic subclone was predicted for each sample, with ETV6.RUNX1.2 having two minor subclones (Supp. Figure 2). We therefore did not observe a correlation between the presence of genetic subclones and observed intra-individual transcriptional heterogeneity, however minor subclones were at the lower end of the allele frequency spectrum and could reflect the neutral evolution frequency tail in some cases^18^. Finally, we looked at somatic mutations at the single cell level using all somatic mutations and retrieved allele calls of individual cells using single cell RNA-seq read alignment files. Around 25% of mutations (n=668) were covered in at least one cell and ∼6.7% (n=185) of them had at least one cell having the mutated allele (Figure 5E). A Fisher’s exact test was run on mutations having at least > 0.1% of cells per sample with the mutated allele (n=31) to identify overrepresented mutations in intra-individual transcriptional clusters. We found somatic mutations in three genes with p-values < 0.1 for samples ETV6.RUNX1.2 and PRE-T2 (intron of C20orf194, 3’ UTR of SH3 domain containing gene *SH3D21* and exon of pyridoxal kinase gene *PDXK*), indicating a slight enrichment of the mutated allele in a transcriptional cluster (Supp. Table 5). Overall very few somatic mutations were fulfilling the criteria for statistical testing and many mutations had a low number of observations contributing to underinflation of p-values and non-significance of all mutations after multiple testing correction (Figure 5F, Supp. Table 5). Thus, it is still possible that other subclonal somatic mutations could be enriched in specific transcriptional clusters but were not possible to detect using this approach. Overall these results suggest that there is some limited evidence linking genetic alterations detected from bulk exome data to intra-individual transcriptional heterogeneity in these samples.

**Figure 5.**
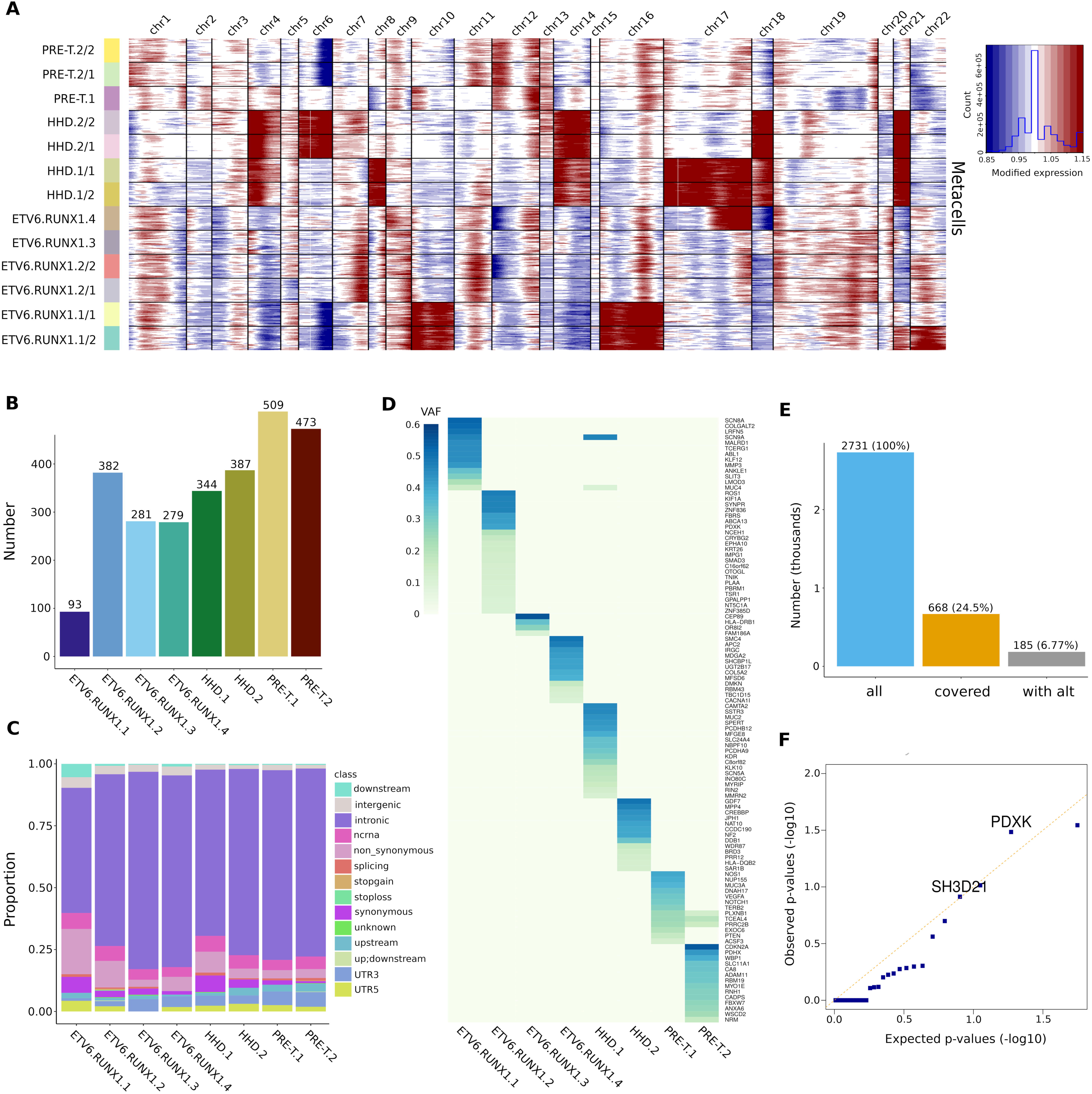
Somatic alterations and intra-individual expression variability. **A)** Copy number profiles of samples and intra-individual transcriptional clusters using equal size metacells and healthy pediatric BMMCs as control cells. **B)** Number of somatic mutations in cALL samples. **C)** Genomic annotations of somatic mutations in cALL samples. **D)** Gene names and variant allele frequencies of exonic non-synonymous somatic mutations in cALL samples. **E)** Number of somatic mutations used as input to obtain allele calls from single cell RNA-seq alignment files using vartrix (<u>all</u> = number of GRCh38 mutations lifted from hg19; <u>covered</u> = number of mutations covered in at least one cell; <u>with alt</u> = number of mutations with at least one mutant allele call). **F)** QQ-plot of Fisher’s exact test p-values of somatic mutations tested for enrichment in intra-individual transcriptional clusters (> 0.1% of cells with mutant allele call, n=31 mutations).

## Discussion

We applied a single cell expression strategy to uncover major sources of intra-individual transcriptional heterogeneity in childhood ALL. To our knowledge this is the first study on intra-individual transcriptional heterogeneity in cALL patients using single cell gene expression. Our study found two notable patterns at diagnosis: first, we found transcriptional heterogeneity associated to the predicted developmental state of leukemia cells which was inversely correlated with the expression of ribosomal protein genes. This observation was more obvious in HHD and pre-T samples but was also seen in in ETV6-RUNX1 samples, indicating a possible subtype specificity. The disrupted products of the fusion between the ETV6 and RUNX1 transcription factors, involved in B cell maturation, could result in an overall lower maturation potential of the leukemia cells. Second, we observed transcriptional heterogeneity linked to gene expression and metabolic regulation, B cell activation and cell proliferation/death in ETV6-RUNX1 samples. We found significant joint over expression of the JUN and FOS proto oncogenes in a cluster of cells for sample ETV6.RUNX1.2, suggesting deregulation in stress/cancer genes.

To further understand this transcriptional variability within samples, we sought to determine if genetic alterations could explain these intra-individual transcriptional clusters. We found no correlation between the number of predicted genetic subclones in a given sample and the presence of intra-individual transcriptional clusters. Indeed, all samples were predicted to have two or more genetic subclones, but not all were found to have intra-individual transcriptional clusters. This could be explained in part by our clustering procedure relying on the ‘most representative’ solution and not returning all major transcriptional clusters. Another explanation could be false positive predictions of genetic subclones triggered by the density of mutations found at the lower allele frequency spectrum and arising from neutral tumor evolution. The low number of somatic mutations in pediatric cancers^19^, coupled with limited coverage of the exome (versus whole-genome) and lower sequencing depth (∼200-300X) compared to targeted sequencing (∼1000-1500X) could have contributed to the problem. We then looked for somatic mutation allele enrichment in specific transcriptional clusters by assigning allele calls to each cell using the single cell RNA-seq read alignment files. Since very few mutations had enough coverage and alternate allele calls to be tested for statistical significance, we failed to report strong evidence of mutant allele enrichment. These results could be explained by the limited expression levels of genes in each cell and the uneven coverage over genes inherent to the 3’ single cell library protocol. We further called copy number variants (CNVs) in each sample to detect events present specific to some transcriptional clusters and found a few large gain and loss events in subsets of cells of ETV6-RUNX1 samples. Given the high number of genes in these regions, the transcriptional impacts of these copy number alterations remain unknown. Taken together, we found some limited evidence of subclonal genetic alterations that were linked to intra-individual transcriptional heterogeneity in cALL samples. As discussed, this could be explained by the resolution of the sequencing data. Generating single cell DNA sequencing data of a targeted set of mutations on these same samples could possibly allow additional discoveries. Relative to this, recent approaches combining scRNA and scDNA data better assigned genetic to transcriptional subclones^20^. Subclonal genetic events altering expression could also be more prevalent in samples where the main source of transcriptional heterogeneity is not related to the predicted developmental state, since maturation potential could be a clonal feature inherent to all cells.

Questions remain about the clinical significance of intra-individual transcriptional heterogeneity in cALL. It is unclear whether subsets of cells at various developmental stages or with variable levels of maturation potential have any fitness advantages during treatment. A previous study looked at the relapse potential of cALL patients at diagnosis using mass cytometry, assigned each individual cancer cell to the closest B cell maturation state and found no correlation between the fraction of cells in a given state and relapse potential^21^. Subsets of cells having deregulated expression of oncogenes (e.g. sample ETV6.RUNX1.2) or genes involved in treatment resistance could be more susceptible to clonal selection. Correcting for developmental state heterogeneity might uncover other subsets of cells having additional relevant transcriptional signatures. Although no patients in our study have relapsed, future single cell studies using matched diagnosis/relapse should provide valuable information on clonal dynamics. Single cell expression data of matched cells should help identify genes and cell surface markers that gain or lose expression in resistant cells and provide insights to guide clinical interventions.

## Methods

### Samples

Study subjects were diagnosed in the Division of Hematology-Oncology at the Sainte-Justine Hospital (Montreal, Canada) and part of the Quebec childhood ALL cohort (QcALL)^22^. Our study cohort consisted of 6 pre-B and of 2 pre-T cALL patients (8 males) with a mean age at diagnosis of 5.8 years and a mean blast percentage of 82% (Table 1). All patients were treated under DFCI protocols. Control subjects were patients for whom the diagnosis of ALL was not confirmed after bone marrow puncture (3 males) and with a mean age of 2.7 years (Table 1). Tumoral DNA specimens were collected from bone marrow mononuclear cells (BMMCs) at initial diagnosis and paired normal DNA specimens were collected from peripheral blood or bone marrow samples without blast cells. All samples were isolated using a Ficoll-Paque gradient fragmentation, washed in PBS and DNA was extracted using the Gentra Puregene Blood Kit from QIAGEN according to manufacturer protocol. For single cells, bone marrow mononuclear cells were cryopreserved in FBS with 10% DMSO. The Sainte-Justine Institutional Review Board approved the research protocols, and informed consents were obtained from all participating individuals and/or their parents.

### Single cell RNA sequencing and analyses

Thawed BMMCs were loaded onto the 10X Genomics Chromium single cell platform (v2 chemistry) at McGill University and Genome Quebec Innovation Center. We aimed for 3,000 cells per sample and targeted 100,000 reads per cell by sequencing each sample on one lane of an Illumina HiSeq 4000 high-throughput sequencer (26v98 b.p. paired-end sequencing) (Table 2). Sequencing reads were processed using the Cell Ranger v2.1.0 pipeline on Ensembl GRCh38.84^11^. BMMCs from three healthy pediatric controls and two publicly available healthy adult control samples (www.10Xgenomics.com) were analyzed jointly using the canonical correlation analysis (CCA) in Seurat v2.3.2^15^. Genes expressed in at least 5 cells, cells having at least 200 genes expressed and cells having less than 8% mitochondrial reads were retained for the analysis. Expression estimates of the genes were regressed out for number of unique molecular identifiers (nUMI), percentage of mitochondrial reads and the S and G2/M cell cycle scores. The thousand most variable genes were selected for each sample and pooled to create a unique list of genes being highly variable in at least two samples (n=883). These genes were kept for the CCA and the top ten canonical correlation vectors were used as input for UMAP v0.2.3^23^. Cell types were identified using the expression of known cell surface marker genes (*CD34/*CD34+, *CD79A/*B cells, *MS4A1/*CD20+ B cells, *CST3/*Monocytes, *GATA1/*Immature Erythrocytes, *HBA1*/Erythrocytes, *CD3D*/T cells, *NKG7*/NK cells, *MZB1*/BCMA). A second analysis was done on cells of three healthy pediatric controls and eight cancer samples. The same cell and gene filters as described above were applied and a Seurat object was created by iteratively merging each dataset. A principal component analysis (PCA) was run on the most variable genes (n=468) and the top twenty principal components were used as input for both UMAP and cluster identification using Louvain’s algorithm^24^ with a resolution of 0.1. Cells from cancer samples that did not cluster with control cell clusters and those that were in the G1 cell cycle phase were retained for downstream analyses. The most variable genes were identified in these filtered cells (n=560) and used as input for both PCA and UMAP. The most representative clustering solution was obtained by finding clusters using resolutions of 0.1 to 3.0 with a step of 0.1. For each of these 30 solutions, we computed all pairwise Adjusted Rand Index (ARI) values and retained the clustering solution with the highest ARI mean (optimal resolution=1.3). Clusters containing less than 10% of cells in a sample were discarded for statistical reasons. Intra-individual differential gene expression analyses were done using the MAST test^25^ (Supp. Table 3). Gene ontology analyses were done using goseq v1.3.0^26^ on the top hundred differentially expressed genes ranked by adjusted p-value. Single cell allele calls of somatic mutations derived from exome data were obtained using vartrix v1.1.0^27^. Copy number profiles of intra-individual transcriptional clusters were obtained by generating thirty equal-sized metacells (sum of raw expression counts of single cells) per cluster that were used as input to infercnv v.0.99.4^28^. Metacells of healthy pediatric BMMCs were used as control. Single cell RNA-seq data are available in GEO under GSE132509.

### Developmental state classifiers and ribosomal protein expression

UMAP coordinates of healthy BMMCs corresponding to the developmental spectrum of B cells (CD34+, B cells and CD20+ B cells) and T cells (CD34+, Immature erythrocytes, T cells) were extracted for input to loess fits. Cells were projected onto the fit line at the closest distance. Outlier cells with distances of more than three standard deviations from the mean were discarded. Cell distances along the fit line were calculated, scaled between 0 and 1 and set as the developmental state pseudotime. Normalized and scaled expression values of the most variable genes determined in the Seurat CCA were extracted and used as input for one-layer fifteen nodes neural networks using the R package nnet v7.3-12. Classifier performances on control cells were assessed using cross-validation of a hundred 70/30 splits and mean RMSE values were computed. Final classifiers were trained on all healthy cells and used to predict the developmental pseudotime of each leukemia cell. Ribosomal protein (RP) expression per cell was defined as the ratio of raw reads mapped to ribosomal protein genes (*RPS-*, RPL-*;* n=97) divided by the total number of raw reads per cell.

### Exome sequencing and analyses

Exome sequencing and analyses were performed as previously described^29^; whole exomes were captured using Agilent’s SureSelect XT Clinical Research Exome kit per manufacturer’s protocol and librairies were sequenced on Illumina high throughput sequencers (HiSeq 2500 or 4000 in paired-end 2×75b.p. or 2×100b.p.) targeting mean coverage of 300X for tumor and 100X for normal. Raw reads were aligned to the hg19 genome reference using bwa v0.7.7^30^, duplicates were marked using Picard v1.107 (http://broadinstitute.github.io/picard/) and indels realigned and bases recalibrated using GATK v3.3.0^31^. Somatic mutations of matched tumor/normal were obtained using mutect v1.1.6^32^, filtered for the PASS flag and variant allele frequency > 0.05 (n=2748, Supp. Table 5). Variants were lifted over using vcf-liftover (https://github.com/liqg/vcf-liftover) for use with single cell RNA-seq data (hg19 → GRCh38, n=2731/2748, 99.4%). Mutations with high coverage (> 100X) were used as input to sciClone v1.1.0^33^ and clonEvol v0.99.11^34^ to generate clonal evolution models. Copy number calls were obtained using sequenza v2.1.0^35^.

## Authors’ contributions

G.B., D.S. and M.C. designed and conceived the study. M.C. and P.S. performed bioinformatic analyses. T.S. and C.R collected and preserved samples. I.R and Y.C.W. supervised and operated the 10X Genomics Chromium single cell platform. M.C., G.B., and D.S. wrote and edited the manuscript.

## Acknowledgments

The authors are indebted to the patients and their parents for participating in this study. Patient tissue samples were provided by the Sainte-Justine UHC Pediatric Cancer Biobank. Computations were run on the Cedar cluster managed by Compute Canada. This work was supported by grants from the FIT program and the Charles-Bruneau Foundation. MC received a PhD studentship from the FRQS. DS holds the research chair Francois-Karl-Viau in pediatric oncogenomics. GB obtained funding from CIHR grant 245250 and is funded by the FRQS.

## Figure legends

<u>**Supplementary Figure 1.**</u> **A)** UMAP representation of cells from three healthy pediatric (n=6,836 cells) and eight cALL (n=32,086 cells) datasets. **B)** Proportion of cells clustering with healthy cell clusters (T + NK cells, B cells + Monocytes, Erythrocytes) per sample. **C)** Cell cycle phases after regressing out S and G2/M phase scores. **D)** Expression of the *CD79A* B cell marker gene in cancer cells. **E)** Expression of the *CD3D* T cell marker gene in cancer cells. **F)** Number of unique molecular indexes (nUMI) in cancer cells.

<u>**Supplementary Figure 2.**</u> Predicted genetic clonal evolution models of cALL.

<u>**Supplementary Figure 3.**</u> Copy number profiles of cALL samples using exome sequencing data.

<u>**Supplementary Figure 4.**</u> B-allele frequency and depth ratio of cALL samples using exome sequencing data.

<u>**Table 1.**</u> Metadata of samples used in the study.

<u>**Table 2.**</u> Single cell statistics of samples used in the study.

<u>**Supplementary Table 3.**</u> Intra-individual differential expression analyses results.

<u>**Supplementary Table 4.**</u> Enriched biological pathways obtained from the top hundred intra-individual differentially expressed genes.

<u>**Supplementary Table 5.**</u> Somatic mutations called from matched tumor/normal whole exome data.

<u>**Supplementary Table 6.**</u> Fisher’s exact test results of mutant allele enrichment in intra-individual transcriptional clusters.

