## Supplementary figures and images for "Single-cell analysis of childhood leukemia reveals a link between developmental states and ribosomal protein expression as a source of intra-individual heterogeneity"

### Supp. Figure 1

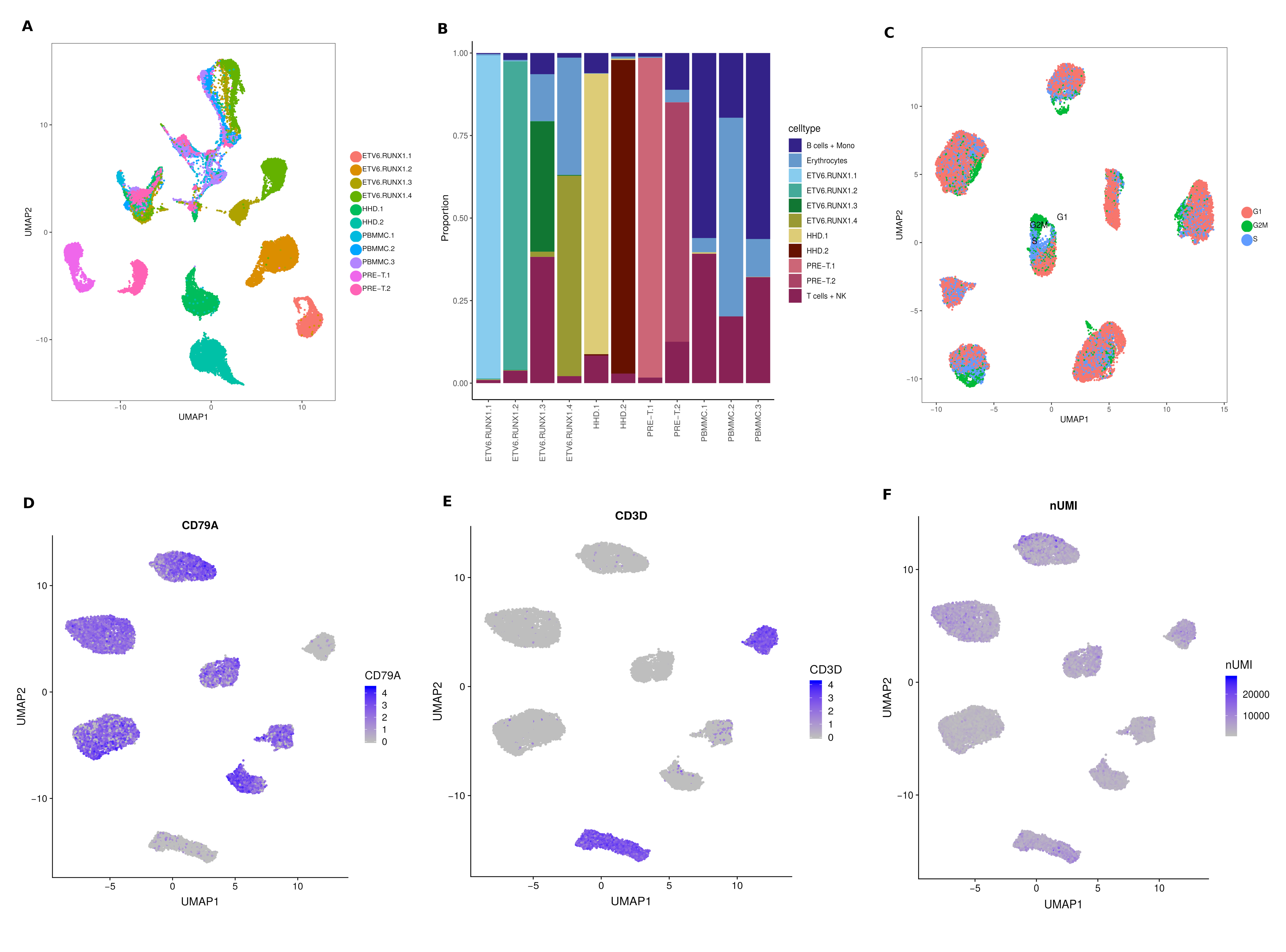

### Supp. Figure 2

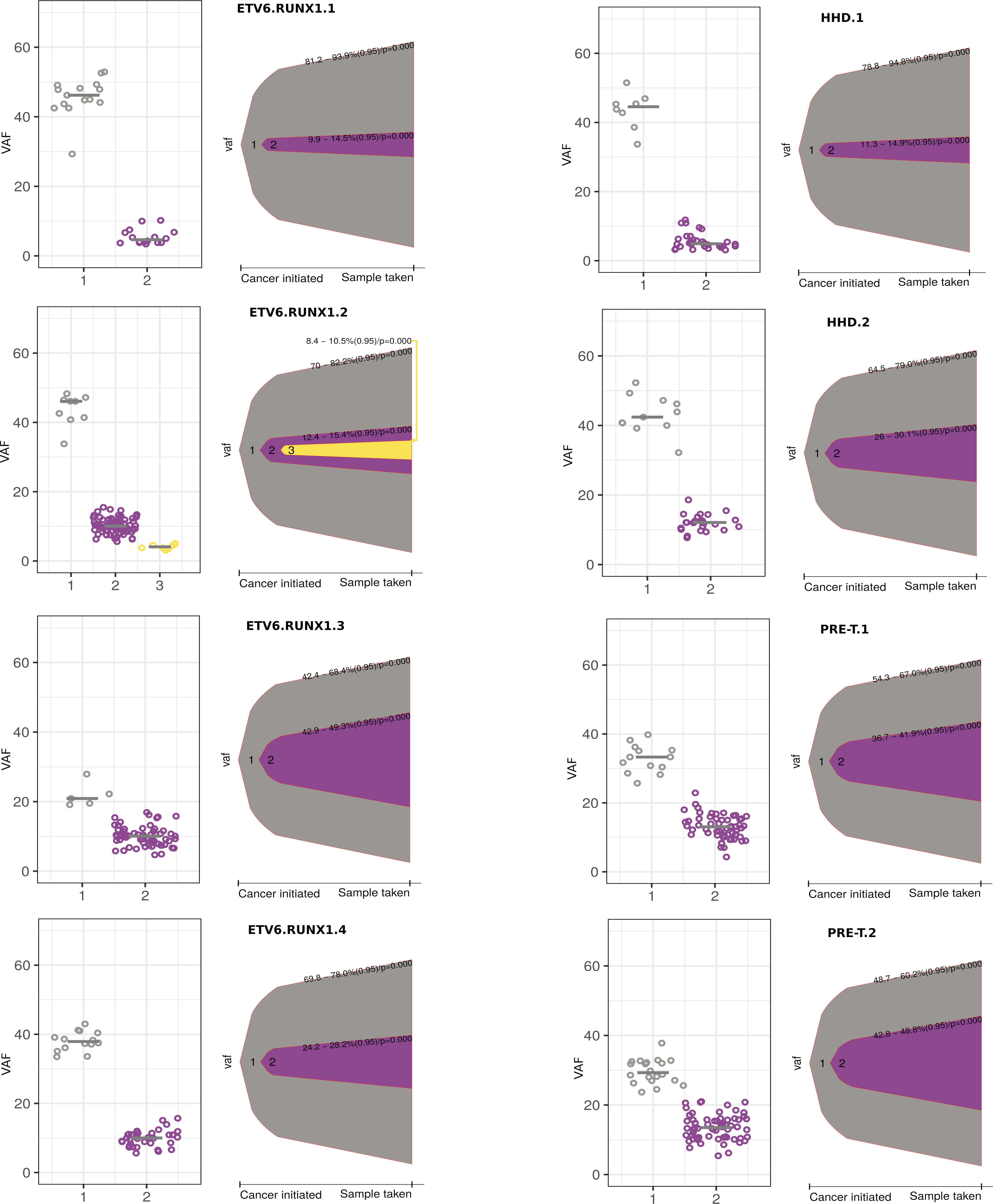

### Supp. Figure 3

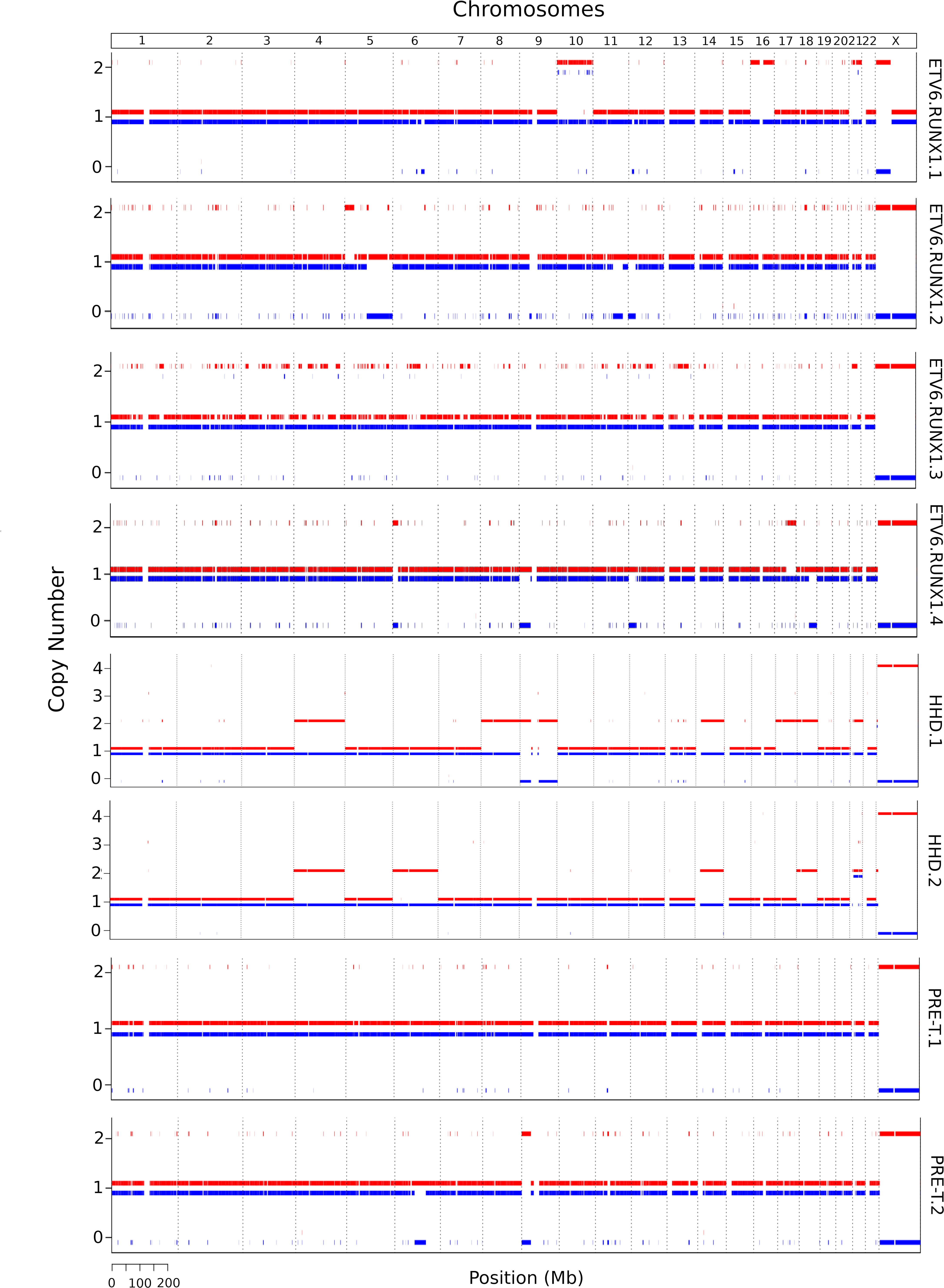

### Supp. Figure 4

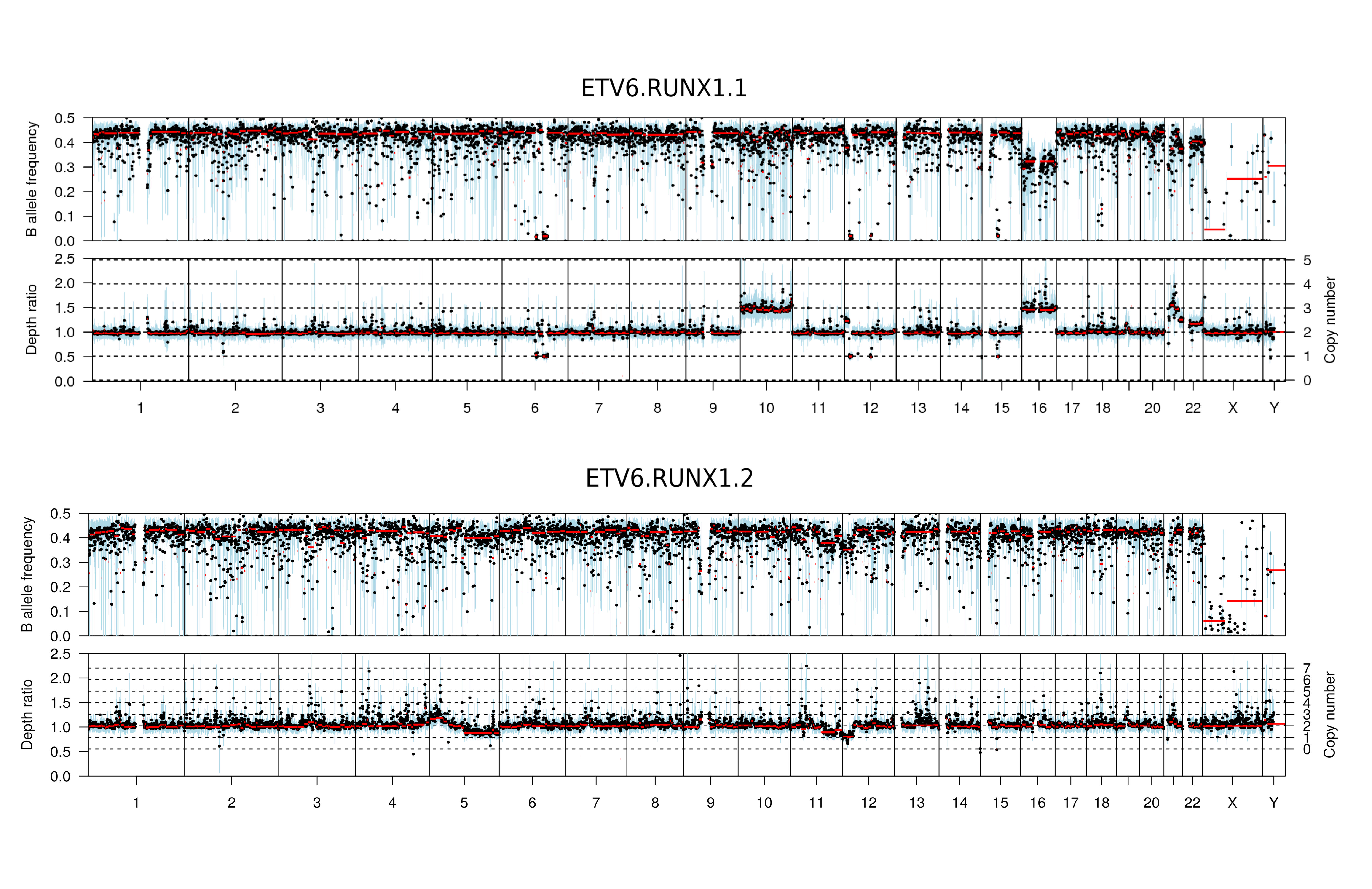
